# Tumor mutational burden shapes success and resistance in cancer immunotherapy

**DOI:** 10.1101/2025.08.09.669487

**Authors:** Guim Aguadé-Gorgorió

## Abstract

Immunotherapy, particularly immune checkpoint blockade, has transformed cancer treatment, yet durable responses remain limited to a subset of patients and cancer types. Many tumors exhibit innate resistance or acquire resistance through immune evasion or neoantigen editing. A central factor in shaping these outcomes is the tumor mutational burden. However, cancer mutations can enhance or impair both cellular replication and immune recognition, reflecting the non-trivial role of mutational load in immunotherapy success and failure. Here, we present a minimal eco-evolutionary model that captures trade-offs between oncogenic and immunogenic mutations in cancer cell replication. Despite its simplicity, the model reveals a rich phase space, including an evolutionary bistable regime where both immunologically silent and mutationally active tumor strategies can coexist. Notably, the model explains two key eco-evolutionary mechanisms of resistance to immunotherapy: preexisting resistance, driven by the persistence of silent clones within genetically unstable tumors; and immunoediting, where immune pressure selects for reduced antigenicity over time.

## I. INTRODUCTION

Immunotherapy has emerged as a powerful approach in cancer treatment, with checkpoint blockade therapies demonstrating remarkable success in a subset of patients [1]. However, therapeutic efficacy remains limited across many cases, and reliable prognostic biomarkers are still lacking [1; 2]. A key factor influencing response is the immunological phenotype of the tumor [3]: *Hot* tumors are characterized by high immune infiltration and neoantigen presentation, making them more likely to respond to therapy, whereas *Cold* tumors show little immune activity and often escape detection altogether, and *Evaded* tumors initially trigger immune responses but evolve towards developing resistance [2]. Understanding what drives these distinct tumor states is essential for predicting immunotherapy outcomes and designing more effective treatments.

Tumor mutational burden (TMB) has emerged as a promising biomarker for immunotherapy prognosis, reflecting the likelihood of neoantigen presentation and subsequent immune recognition of malignancies [4; 5]. However, its predictive power remains inconsistent: high TMB does not guarantee therapeutic success [6; 7], and neoantigen heterogeneity [8], lack of immune infiltration [5] or the evolution of escape mutations can all lead to immunotherapy failure [2]. High TMB can drive both heightened immune recognition and, paradoxically, strong selection for immune escape mutations [2; 9]. Conversely, low to intermediate TMB tumors may preferentially evolve through immunoediting, reducing antigen presentation via HLA loss or epigenetic silencing [9; 10; 11]. To understand the nuances of how mutational load and genome instability determine tumor immunogenicity, we must acknowledge that these events are driven by ecological and evolutionary processes [12]. Mathematical models of cancer eco-evolution and immunogenicity provide a necessary framework to explore immunotherapy resistance [13; 14; 15].

Recent theoretical work has begun to explore the role of mutational dynamics in cancer-immune interactions. In particular, mathematical models validated with experimental or clinical data have corroborated the role of genome instability [16], neoantigen heterogeneity [17], or immunoediting dynamics [10] in shaping cancer immunogenicity. Yet, it remains unclear whether a tumor will respond to immunotherapy, remain cold and immunologically silent, or else will evolve immunotherapy resistance by silencing genes or activating escape mutations. We still lack a unified conceptual framework explaining the role of TMB in driving each of these evolutionarily trajectories.

In this work, we study a minimal eco-evolutionary framework based on an evolutionary landscape in which cancer cell replication depends on the fraction of the cell genome that is mutated (Fig. 1). Previous research proposed that cancer cell replication can be modeled to increase with oncogenic mutations, provided no essential housekeeping genes are compromised [18; 19]:

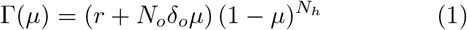

**FIG. 1.**
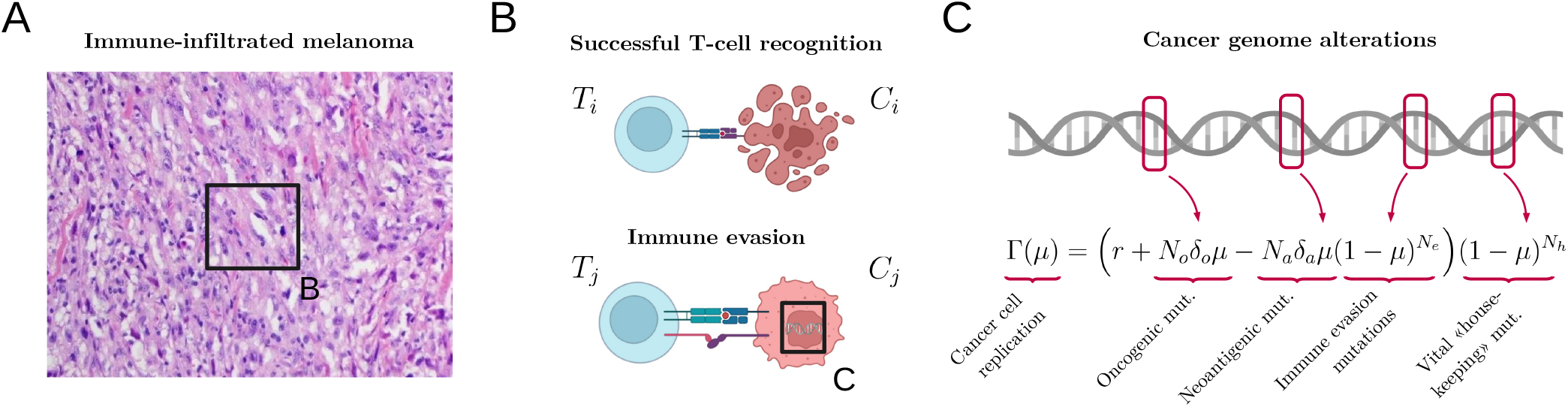
Impacts of tumor mutational load on cancer cell replication and survival. (A) Highly mutational tumors like melanoma are often infiltrated by immune cells and treated with immunotherapy, yet with variable responses (Image from Wikimedia Commons, with license CC-BY-SA-3.0). (B) A complex set of conditions controls whether cytotoxic T-cells are able to detect and kill tumor cells (top), or else these find mechanisms to avoid immune recognition (bottom). (C) A key determinant of T-cell success is the mutational load of a given cancer cell (*µ*). Yet, the impact of mutations on cell replication and immunogenicity is multidimensional, here captured by a replication function Γ(*µ*) that increases or decreases depending on the type of mutations being activated.

Here, Γ is the replication rate of a given cell and *µ* represents the fraction of mutated genes in its genome, serving as a proxy for the effective mutational load at a given time and hence the relative TMB [19; 20]. *N*_*o*_, *N*_*h*_ are the number of oncogenic and house-keeping genes respectively (see Materials and Methods), and *δ*_*o*_ defines the average effect of oncogenic mutations on replication. The Γ(µ) function therefore describes a minimal evolutionary landscape that links mutations to replication via a trade-off: replication is enhanced by mutations in oncogenes, but is rapidly impaired if any of the *N*_*h*_ house-keeping genes are mutated. This trade-off predicts the emergence of unstable, fast-replicating clones as an optimal evolutionary strategy, consistent with the biology of microsatellite-unstable tumors [20; 21].

However, this landscape omits the critical relation between mutational loads and immune surveillance in tumors [22; 23]. In particular, CD8^+^ T cells can recognize and eliminate cancer cells through detection of neoantigenic mutations, a key mechanism in major immunotherapy strategies [22]. Other mutations, conversely, may confer resistance to immune surveillance [2] (see Materials and Methods). In sum, the TMB plays a non-trivial role on cancer cell replication, mediated by two central trade-offs in both cell replication and immunogenicity. To address the complex role of the TMB in shaping immunotherapy success or failure, we extend the replication landscape above in order to include the effects of both immunogenic and immune-evasive mutations. This provides a minimal framework to explore how mutational processes drive tumor immunogenicity and resistance to immunotherapy.

## II. RESULTS

### A. A minimal eco-evolutionary trade-off generates four tumor immunogenicity regimes

To include the ecological pressure of immune surveillance we study a trade-off between mutations that lead to immunogenic antigens (and hence immune detection and cell death) and mutations that allow for immune evasion and reduce immune killing [2; 22]. To do so, we can define immune-mediated death as a function of the fraction of the mutated genome as

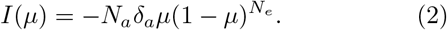

Here, *N*_*a*_ represent the number of genes that result in neoantigenic petides, and *δ*_*a*_ the average effect of these neoantigens in immune recognition [24]. Mutations in neoantigens decrease cell viability following a linear decay as estimated in [16; 17], although empirically-inferred non-linear functions could also be implemented (see e.g. [10]). Additionally, mutations in immune-escape genes *N*_*e*_ can decrease the effect of immune recognition, and the full trade-off implies that TMB decreases cell viability unless immune escape mutations are activate. The complete replication landscape is now

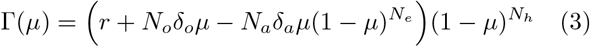

The first question is to understand how *µ* and the oncogenic-immunogenic state {*δ*_*o*_, *δ*_*a*_} modulate the shape of this function and the presence of maxima, representing optimal TMB states for cancer cell growth. We find that Γ(*µ*) unfolds a rich and interpretable ecoevolutionary landscape capturing the complex role of the TMB in cancer immunogenicity (Fig. 2A). The phase space contains four distinct immunogenic regimes, each corresponding to qualitatively different states of the cancer-immune interaction separated by analytically predictable critical lines (see Materials and Methods for all numerical and mathematical details).

**FIG. 2.**
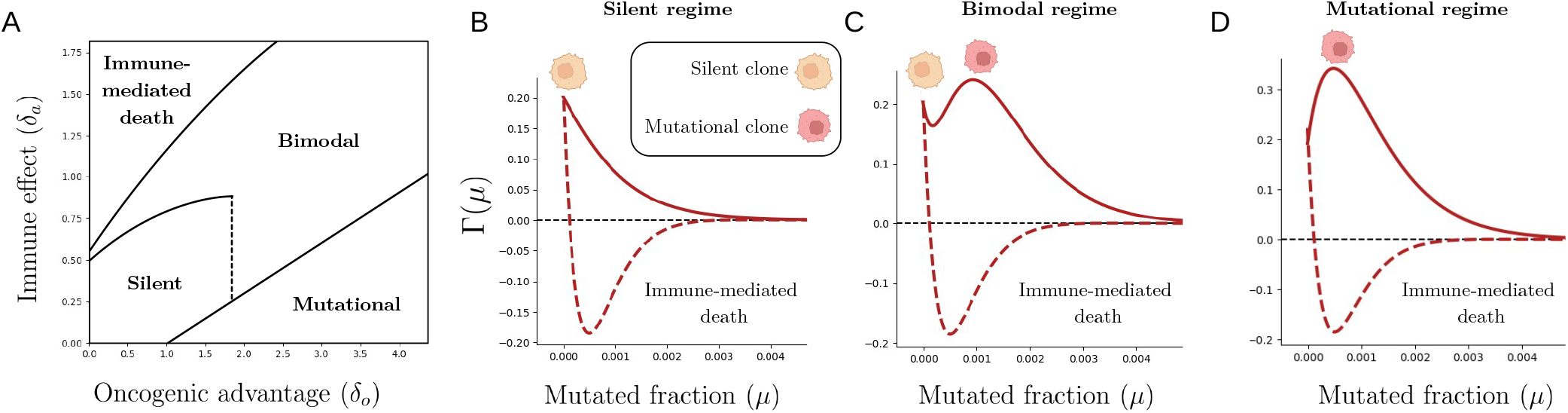
Phase space of the possible immunogenicity regimes. (A) The model unveils a phase space with four possible configurations of the replication function Γ(*µ*) depending on the effect of oncogenic (*δ*_*o*_) and neoantigenic (*δ*_*a*_) mutations. (B) In the absence of a strong immune infiltration or mutational advantage, tumor cells remain silent. Therapeutic increase of *δ*_*a*_ (dashed lines) will not result in cell death. (C) When *δ*_*a*_ and *δ*_*o*_ are balanced, the landscape reveals a bimodal state in which both silent and mutational populations are optimal replicators. Therapeutic increase of *δ*_*a*_ might eradicate the mutational clones, allowing competitive release of silent clones. (D) When *δ*_*a*_ is very low (e.g. lack of infiltration or activated immune escape), tumors are pushed towards instability and high mutational burdens. Therapeutic recovery of *δ*_*a*_ might result in tumor arrest, but also induces selective pressures towards immunoediting.

#### Silent regime

When both immune and oncogenic effects are weak, there is little selective pressure for accumulating mutations. The replication landscape remains a decaying function, favoring silent, stable phenotypes without high mutational burdens (Fig. 2B). This scenario aligns with cancers that arise via epigenetic dysregulation and exhibit minimal mutational burdens, such as early-onset and epigenetically driven pediatric leukemias like acute myeloid and lymphoblastic leukemias [25].

#### Mutational regime

If oncogenic benefits are strong and immune pressure is weak, the landscape favors the progression towards a single optima with mutated cancer cells (Fig. 2D). In it, tumors evolve toward high mutated genome fractions, neoantigen presentation and overall genome instability. This regime includes tumors with high TMB and pervasive genomic instability that are often sensitive to checkpoint blockade, such as cutaneous melanoma, smoking-associated nonâsmall cell lung cancer, and microsatellite-unstable colorectal or endometrial carcinoma [8; 23].

#### Bimodal regime

When oncogenic benefit or immune pressure are not dominant, the function exhibits two different maxima (Fig. 2C, see Materials and Methods): both silent and mutationally active phenotypes are locally optimal. This key regime implies that the same tumor microenvironment can support coexisting strategies: either remaining genomically silent or evolving mutationally via immune evasion result in increased replication. Heterogeneous immunogenicity has been reported in the high-TMB cancer types described above [8], as well as in triple-negative breast cancers and renal cell carcinomas with evidence of coexisting hot and cold regions within the same tumor [26; 27]. This result provides a theoretical framework to understand the presence of silent, immune-resistant clones inside aggressive and unstable tumors (see next section).

#### Immune-mediated death regime

When immune pres-sure is strong, such as after effective checkpoint blockade inhibition or engineered interventions like CAR-T therapy, the cost of immunogenicity outweighs oncogenic benefit. In this regime, mutationally active clones are strongly suppressed, reflecting the deep tumor regressions observed in responders to checkpoint blockade [23]. However, silent clones with minimal neoantigen presentation often persist, and robust immune elimination imposes a selective gradient favoring immune-silent states: survival will always be granted for low enough *µ* even if therapy ensures strong *δ*_*a*_ (Fig. 2). The fact that low TMB states still have high Γ values under strong immune attack explains clinical observations where tumors initially respond dramatically, but later relapse due to evolution of antigen silencing [11; 28], as studied in section II.C.

Together, these regimes show that different tissue and microenvironmental constrains, modulating *δ*_*o*_ and *δ*_*a*_, can lead to different scenarios in which TMB modulates cell replication and immunogenicity differently. Importantly, the bimodal regime implies that mutational and silent clone heterogeneity is an intrinsic property of the eco-evolutionary process at play, not just a transient feature. One possible exploration at this stage would be to perform evolutionary dynamics on top of these complex landscapes throughout the full parameter domain. Instead, we focus on two specific scenarios that are important for immunotherapy resistance: (i) Under what conditions can silent and mutational clones stably coexist within a tumor, and (ii) What factors determine whether high-TMB tumors are eradicated, or else evolve resistance through immune escape or antigen silencing.

### B. Preexisting resistance: silent clones inside immunogenic tumors

The two mutational trade-offs in Γ give rise to a regime with two distinct replication optima, separated by a valley with lower (yet still positive) replication (Fig. 2C). The peaks correspond to a mutational phenotype and a silent phenotype with lower immunogenicity. The presence of two local optima in replication indicates that these might be stable evolutionary outcomes shaped by opposing pressures on growth and immune visibility.

We analyze the endpoints of evolutionary trajectories in which the evolved trait is the fraction of expressed mutations *µ* (see Materials and Methods). The silent phenotype at *µ* = 0 represents an absorving state of the evolutionary dynamics (not a true evolutionary stable strategy, ESS), while the mutational optimum is an ESS [29]. Additionally, both states are separated by a branching point at intermediate TMB [30]. Under this configuration, silent clones would remain around *µ ≈* 0, unless large enough mutational events such as loss of mismatch repair [31] push some cells beyond the branching point and towards the mutational state.

This raises the question of whether the two phenotypes can coexist inside a single tumor. Classical ecological models suggest they cannot: in a well-mixed system described by the replicator equation, the faster replicator outcompetes the other, leading to classical results of competitive exclusion (see Materials and Methods). In the presence of genome instability, the replicator-mutator model shows that the mutational clone could maintain an immunoge-silent population in place by generating mutants with much lower immunogenicity (see Materials and Methods). Even if silencing all mutational signatures (*µ ≈* 0) is virtually impossible in cancer [7; 20], the model predicts that the mutator phenotype could maintain at least a population that has lost part of its immunogenicity even when silencing events are extremely rare (see Materials and Methods and next section).

Beyond the rare possibility of silencing events, real tumors are spatially structured, which is known to alter the outcomes of competitive exclusion [32]. To explore this, we develop simple simulations of stochastic birthdeath dynamics on a 2D grid inspired by models of drug sensitive-drug resistant tumors [33]. We simulate random and spatially structured tumors, in which the slowest replicator is located approximately inside a disc at the center of the grid. Additionally, we consider well-mixed and first-neighbour replication rules (see Materials and Methods), and show in figure 3C the mean and distribution of time to extinction of the slowest replicator. We find that, although the mutational strategy eventually dominates, clonal clustering and local replication greatly extend the persistence of the slower clone. Technically, local replication is a necessary condition for increased survival (as opposed to mean-field replication), while clonal clustering further enhances this effect (Fig. 3C). Spatial constrains create refugia where silent clones can survive, even within rapidly growing mutational tumors, and potentially re-emerge after therapy. These effects are consistent with empirical observations of spatial structure preserving intra-tumor diversity [32; 34; 35] as well as distinct immune-hot and cold spatial domains over long timescales [26; 27].

**FIG. 3.**
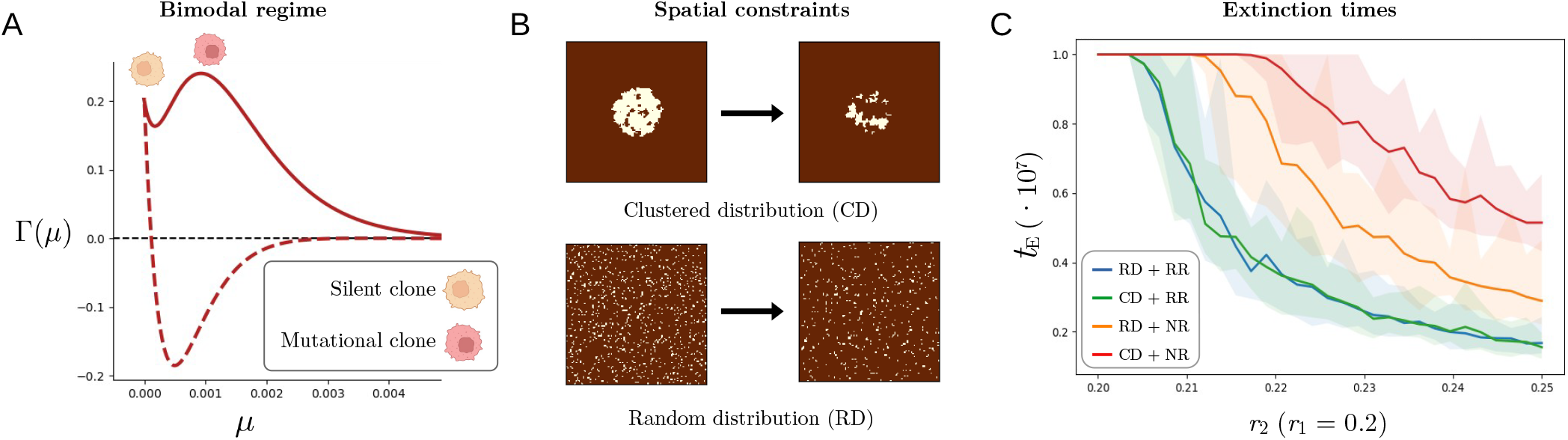
Silent clones can survive inside immunogenic tumors. (A) Oncogenic-immunogenic trade-offs predict a regime in which both silent and mutational strategies are replication optima, yet mean-field models indicate that the fastest replicator would exclude slower populations. (B) Different spatial constraints can modulate the time to extinction. Simulations of clustered (top, CD) and random (bottom, RD) initial spatial distributions, as well as neighbor (NR) and random (RR) replication schemes (see Materials and Methods). (C) Mean (lines) and distribution (shaded areas) for the times to extinction of a mutationally silent clone *c*_1_ (*r*_1_ = 0.2), given stochastic spatial competition with a mutationally active, faster replicator clone *c*_2_ with increasing *r*_2_ *r*_1_ (see Materials and Methods for details of the simulations). Both local replication and clustered clone distributions slow down the time to extinction, allowing for multi-clonal coexistence.

These results suggest that evolutionary bimodality and spatial structure enable prolonged coexistence of im-munologically distinct clones. This puts immunotherapy within the context of Adaptive Therapy (AT) [33; 36]: in AT, drug scheduling avoids a maximum dosage treatment that would kill all drug-sensitive cells and lead to a competitive release of drug-resistant clones. Instead, the aim of AT is to disrupt tumor growth by maintaining a certain drug-sensitive population in place. Our work motivates the possibility of a new *Adaptive Immunotherapy* : if immunologically-silent clones struggle to survive inside highly immunogenic tumors (Fig. 3C), therapy scheduling of CAR-T dosage could maintain intratumor clonal competition to avoid the relapse of a silent clone that is fully resistant to immunotherapy [37]. Effective immunotherapy must consider the underlying spatial ecology of weakly-immunogenic states and clonal competition.

### C. Acquired resistance: immune escape vs neoantigen editing

In the first formulation of the model, all mutations are aggregated into a single relative TMB trait *µ*, which is useful for building intuition about mutational trade-offs, but obscures important mechanistic distinctions. In reality, selective pressures can lead to survival of cells with different mutational burdens across mutation classes: immune surveillance might select for cells with a lower neoantigen load (lower *µ*_*a*_) or increased escape mutations (higher *µ*_*e*_). These correspond to two well-documented resistance mechanisms: (i) *immunoediting*, whereby tumors reduce or lose antigen expression [2; 11], and (ii) ac-quisition of escape mutations that actively block immune killing, such as PD-1/PD-L1 or CTLA-4 pathway alterations [2; 28; 38]. The therapeutic implications are profound: checkpoint blockade can reverse escape-mediated resistance, but it is ineffective against fully immunoedited clones [28].

To understand the evolutionary mechanisms driving either editing or escape, we rewrite the immune surveillance function taking into account the fraction of mutations affecting neoantigenic and immune escape genes separately:

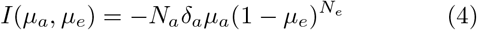

Both lowering *µ*_*a*_ or increasing *µ*_*e*_ reduce immunemediated death. If *µ*_*a*_ *≈* 0, there is no immune recognition and no need for *µ*_*e*_. If *µ*_*e*_ is high, immune cytotoxicity is blocked and neoantigen loads *µ*_*a*_ no longer lead to immune elimination (Fig. 4A). Which strategy is favoured under a given eco-evolutionary setting can be captured by the selection gradients, which in the simplest scenario are equivalent to the gradients of the replication function (see Materials and Methods):

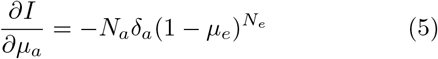

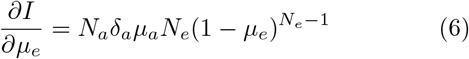

**FIG. 4.**
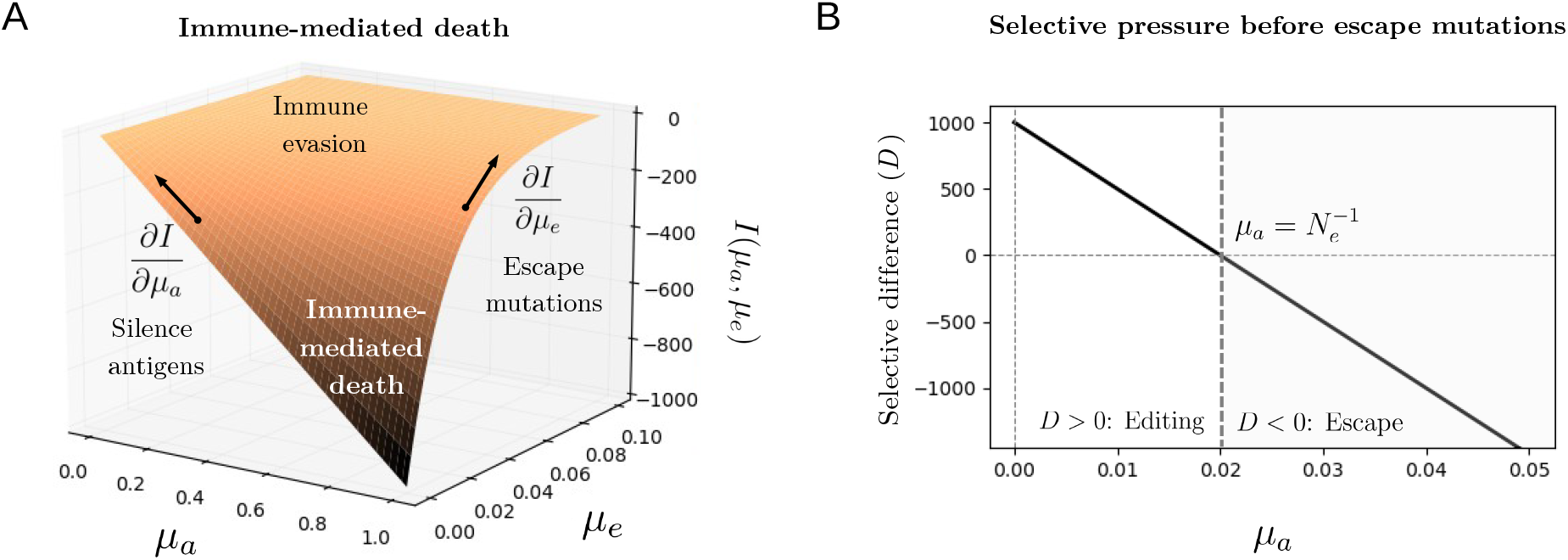
Immunotherapy resistance follows immune escape or antigen editing pathways. (A) Immune surveillance can be attenuated by cancer cells by either reducing the fraction of neoantigenic mutations (immunoediting, *µ*_*a*_ ↓) or increasing the fraction of immune-escape mutations (*µ*_*e*_ ↑). Which strategy is most benefitial depends on the gradients of the landscape, which in turn depend on the current immunogenic state *µ*_*a*_, *µ*_*e*_. (B) Before evasion mutations are active (*µ*_*e*_ ≈ 0), the model predicts that, at low *µ*_*a*_, selection for antigen silencing is stronger, whereas at higher *µ*_*a*_ escape mutations provide a higher evolutionary advantage. This is consistent with clinical data showing that immune escape is favoured in tumors with higher TMB [9] or pervasive genome instability [10], while weakly or moderately antigenic tumors are typically subject to antigen editing [10].

The sign and magnitude of these gradients determine whether silencing (*µ*_*a*_ ↓) or escape (*µ*_*e*_ ↑) is favored (Fig.4A). This depends on the mutational state of the tumor, and can be captured by the difference *D* between absolute selection pressures

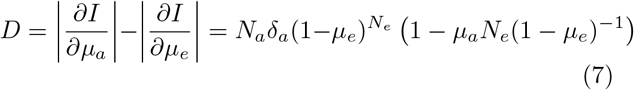

A positive *D* value indicates selection of immunoediting, while a negative *D* value indicates selection of immune-escape mutations. A key question is to understand which evolutionary mechanism dominates at initial stages of tumor growth. Before escape mutations are active (*µ*_*e*_ *≈* 0), we find that the difference in selective pressures simplifies to (Fig. 4B):

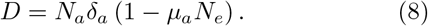

Before escape mutations are active, the favoured strategy depends on the relative neoantigenic load *µ*_*a*_ (Fig. 4B). If there are few antigenic mutations, selective pressures for immunoediting are stronger. Instead, if the relative neoantigen load is larger than 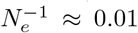, cor-responding to about 30-50 antigens (see Materials and Methods), silencing antigens is no longer a strong-enough strategy, and escape mutations will be selected by immune pressure. This suggests that, at initial stages of tumor growth, whether immune surveillance favours immunoediting or immune escape depends on the current antigenicity of the tumor.

This rule-of-thumb result is consistent with clinical evidence of immunotherapy resistance. In particular, recent research has shown that escape mutations (like checkpoints PD-1/PD-L1 or CTLA-4) are most commonly found in tumors with high microsatellite instabillity [10] or high TMB [9], while tumors with moderate TMB values and antigen counts are more likely subject to immunoediting [**?**]. Overall, the simple immunogenicity landscape of figure 4 provides a conceptual framework to understand how TMB modulates immunotherapy resistance at different stages of tumor growth.

## III. DISCUSSION

Immunotherapy has transformed cancer treatment, yet resistance remains a major challenge. Resistance to immunotherapy and T-cell recognition manifests in diverse forms, from primary resistance, where tumors are immunologically “cold” and unresponsive from the outset, to adaptive resistance, where initially responsive tumors relapse through the evolution of immune escape or neoantigen silencing [2; 11]. A core finding in immunotherapy is that tumor mutational burden (TMB) plays a paradoxical role in immunotherapy outcomes. While high TMB is often associated with improved checkpoint blockade response [39; 40], clinical outcomes remain inconsistent, and a clear mechanistic framework is lacking.

In this work, we have studied a minimal and conceptual eco-evolutionary model to explore how TMB shapes immunotherapy outcomes. While the model omits fundamental features in cancer-immune interactions such as stromal suppression [41], T-cell exhaustion [42], or myeloid-derived pro-tumor immunity [43], it formalizes a core trade-off between antigen load and immune escape, which remains a primary axis of resistance evolution. A central outcome of the model is that, only by imposing simple trade-offs between different types of mutations, the model already reveals a surprisingly rich landscape of tumor immunogenicity states. In particular, the landscape captures the existence of cold tumors, where cells remain silent and mutations are suppressed, and the presence of a regime where most immunogenic cells are recognized and killed by the immune system.

In between these two extremes, the model uncovers simple rules behind two key immunotherapy resistance mechanisms. First, it highlights the mechanisms by which silent and mutationally-active clones can coexist inside a tumor, highlighting the need to revisit current immunotherapies with the principles of adaptive therapy and spatial heterogeneity in mind [33; 34; 36]. Second, the model uncovers an evolutionary dichotomy between immune evasion via mutational escape or via antigen silencing that is consistent with clinical observations, in which high TMBs are linked to escape mutations, while low or intermediate TMBs may favor antigen silencing and immunotherapy resistance [9; 10].

This work underscores the value of eco-evolutionary thinking in cancer immunology. By abstracting complex dynamics into interpretable and qualitative principles, we uncover simple rules governing resistance pathways and their dependence on tumor mutational burden. Integrating more advanced versions of this eco-evolutionary model with the ecology of T cell dynamics and patientspecific TMB data may enable rational, adaptive scheduling of immunotherapy that exploits evolutionary tradeoffs to sustain tumor control.

## ACKNOWLEDGMENTS

Special thanks to Ricard Solé, with whom many of these ideas have been discussed throughout the years, and to Sonia Kéfi, Victor Maull and Jordi Piñero for their support. The author is supported by a Marie Sk-lodowska-Curie Actions Postdoctoral Fellowship under project FRAGILEPRINTS - 101105029. Views and opinions expressed are however those of the author(s) only and do not necessarily reflect those of the European Union or the CNRS. Neither the European Union nor the CNRS can be held responsible for them.

## MATERIALS AND METHODS

### On the number of genes

In our model, the replication rate Γ(*µ*) depends on a set of parameters *{N*_*i*_*}* that represent the number of genes belonging to functionally distinct families relevant to cancer evolution. The number of such genes can vary from cancer type and even patient to patient, and here they only represent a proxy for the dimensionality of the eco-evolutionary processes in place. This means that our model does not seek to use the exact value for each *N*_*i*_. Instead, and in line with the qualitative goal of this project, we provide approximate numbers for the orders of magnitude of the gene family sizes. Quantitative changes can lead to different areas in the phase space, but the qualitative presence of 4 general regimes remains in place. Additionally, different genes can have extremely diverse lengths in terms of base pairs [44], so that mutations would not necessarily be equiprobable across gene types. In this simple approximation, the four gene families are defined as follows: *N*_*o*_: Oncogenic driver genes. This parameter denotes the number of genes whose mutations confer a selective advantage via increased proliferation or survival (e.g., KRAS, MYC, PIK3CA) [45]. nalyses of cancer genomes suggest 100-300 genes commonly function as oncogenic drivers [45; 46]. This number has a strong dependency on the cancer and tissue type and microenvironmental properties in place (larger *N*_*o*_ leads to a larger single-peaked domain, as expected). Across the paper we use *N*_*o*_ = 300 to depict a highly-oncogenic tumor. *N*_*a*_: Immunogenic (neoantigen-generating) genes. This represents the number of genes whose mutations can result in neoantigen formation, making the tumor visible to the adaptive immune system [22]. As most protein-coding genes can, in principle, generate neoantigens, this family spans a broad range. However, not all expressed mutations are presented or immunogenic, and evidence indicates that many neoantigenic mutations lead to weakly-detectable antigens, whereas fewer might generate strong-binding antigens [24; 47]. Assuming approximately 20.000 protein-coding genes in the human genome, accumulated evidence indicates that as many as 15-20% of the whole genome could yield immunogenic neoantigens [24; 48; 49; 50], and we set as an example *N*_*a*_ = 3000 mutations that can be processed and presented effectively. Increasing this number leads to an increase of the immunemediated death domain in the phase space. *N*_*e*_: Immune evasion genes. These are genes whose loss or alteration reduces immune detection, including MHC class I/II genes, antigen presentation machinery (e.g., B2M, TAP1/2), checkpoint regulators (PD-L1, CTLA4 pathways), and IFN-*γ* response genes [2; 51]. Based on pathway analyses and CRISPR screens, 50-100 genes are frequently implicated in immune evasion [52; 53; 54]. We set this number to be *N*_*e*_ = 100. *N*_*h*_: Housekeeping / essential genes. This refers to genes critical for basic cellular function, where mutations result in lethality or loss of viability. Large-scale screens like [55; 56; 57] identify 10-15% of human genes as essential across cell types. Out of 20,000 protein-coding genes, we set here *N*_*h*_ = 2500. These parameter estimates are order-of-magnitude approximations, intended to capture the minimal trade-offs between proliferation, immunogenicity, immune escape and cellular viability under mutational pressure. Exact values may vary by tissue, tumor type, and immune context.

### Analytical predictions of the phase space boundaries

In figure 2A we provide a phase-space for the different immunogenicity regimes, in which the critical lines separating phases are provided by the following analytical and semianalytical techniques. The first line at the bottom of the phase space separates the single-peaked regime dominated by a mutational tumor. To observe this peak, the requirement is that the first mutations are already benefitial, so that the landscape starts with an uphill gradient leading to evolution-ary increases of the mutational load. This translates to finding a positive gradient of Γ at *µ* = 0, so that the critical line is found when the gradient is null:

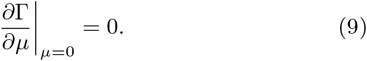

Because this derivative will be studied several times, it is worth writing it in detail. If we re-name

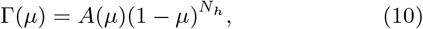

the derivative writes

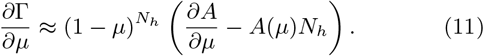

The sign comes from the fact that we have used 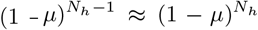 after the derivation, which is a good approximation for large *N*_*h*_ values (*N*_*h*_ *≈* 2500). By derivating *A*, we find

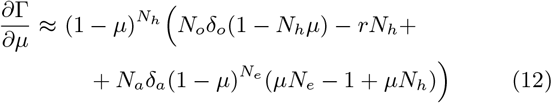

At *µ* = 0, the signature for a mutationally-inactive clone, the gradient becomes

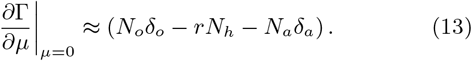

The critical line separating the mutational tumor from the silent or bimodal states is found at

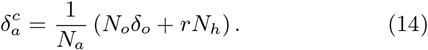

If the effect of immune recognition and attact *δ*_*a*_ is below this threshold, the gradient at *µ* = 0 will be positive and any silent state will have an evolutionary pressure towards accumulating mutations (Mutational regime).

The second line separates a regime with no peaks, where Γ(*µ*) only decreases (silent clone), from a regime with two peaks, in which Γ(*µ*) becomes bimodal and separates an optimal at *µ* = 0 (which is not necessarily a peak in the sense of an Evolutionary Stable Strategy, but an optimal as lower *µ <* 0 values are unrealistic) from an optimal at higher mutational loads. This transition is marked by the emergence of two points in the *µ >* 0 domain in which Γ(*µ*) becomes flat, indicating local minima and maxima. This is found by equating *∂*Γ*/∂µ* to zero. Using the approximation developed above, the equation to solve reads:

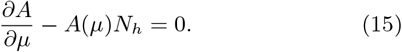

Yet, from (11) we can see that this equation cannot be solved exactly. Instead, we can simplify the numerical search for the conditions under which peaks and valleys emerge. In the area close to *δ*_*o*_ *≈* 0, *δ*_*a*_ *≈* 0 (the silent state regime), we know that *∂*Γ*/∂µ* is uniformly negative across the *µ >* 0 domain. Thus, the emergence of peak-and-valley can be translated to the emergence of a parameter domain (*δ*_*o*_, *δ*_*a*_) in which the gradient is negative at *µ* = 0,

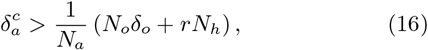

yet it recovers after a valley and becomes positive again:

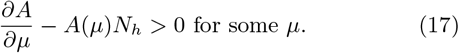

We implement (15)-(16) numerically by searching, for each value of *δ*_*o*_, for the first *δ*_*a*_ value that fulfills both conditions. The immune death regime is characterized by a domain in which Γ(*µ*) becomes negative, knowing that Γ is positive at *µ* = 0 and tends to 0^+^ at *µ* → 1. We perform a similar numerical search for the first *δ*_*a*_ value given *δ*_*o*_ at which

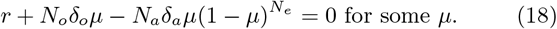

### Replicator-mutator equations and multiclonal coexistence

In our model we study the simple ecology of the relative abundances of two competing cancer populations, *c*_*s*_ (silent) and *c*_*µ*_ (mutational), competing for space within the context of a solid tumor (Fig. 3). Because the only cell trait is the replication rate defined by Γ(*µ*) and we do not have information of the cell-cell interactions that would define a Lotka-Volterra cancer model [15], the Replicator equation is the best-adapted model to describe our system [58; 59]. We study the ecology of relative abundances similar to a constant population constraint (*c*_*s*_ + *c*_*µ*_ = 1) representing a dense solid tumor, and refer the reader to classical literature also discussing replicators in growing populations [60]. In this context, the equations governing mean-field dynamics constrained by spatial limitations are

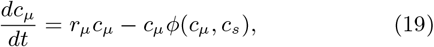

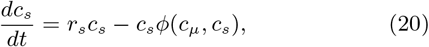

where *ϕ* = *r*_*µ*_*c*_*µ*_ + *r*_*s*_*c*_*s*_ to fulfill the constant population constrain. Writing *c*_*s*_ = 1 − *c*_*µ*_, we arrive to

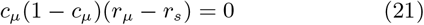

as the condition for the equilibrium state of the *c*_*µ*_ population. If *r*_*s*_ *> r*_*µ*_, *c*_*µ*_ = 0 is the only stable attractor, and if *r*_*µ*_ *> r*_*s*_, *c*_*µ*_ = 1. Thus, the fastest replicator will eventually populate all space as typically predicted by the competitive exclusion principle [61].

Another key mechanism that can impact the outcome of the replicator equation is the possibility of mutational shifts between cancer populations. As a toy model to capture the meaning of those dynamics, we assume a system in which the *c*_*µ*_ population is characterized by genome instability. In this context, cells from the *c*_*µ*_ population, by silencing pieces of their genome, can transition towards an approximately silent 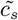 state. Although this is an extreme assumption, because silencing all the mutations in a cancer genome is virtually impossible, it sets the stage for the dynamics described in the last section of the Results, in which cells can silence, at least, those mutations related to neoantigen presentation in the presence of immune surveillance. In this context, and assuming that what we label as 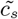 now is not a fully silent (*µ ≈* 0) state, but rather a cell that has lost most of its immunogenicity, on could think of a Replicator-Mutator ecological process following [59]:

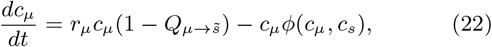

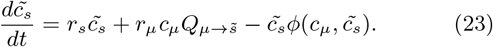

In this setting, we are defining the mutational shift 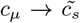 with rate *Q* as a process happening during replication, indicative of replicating errors related e.g. to mismatch-repair defficiencies. Alternative models that study phenotypic plasticity and shifts unrelated to replication reach conceptually similar results to those exposed here [62; 63]. Writing again 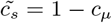, the first equation becomes

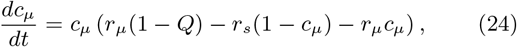

from where a stable abundance for a coexistence state is possible at

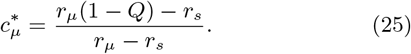

To ensure that 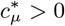 in the assumption of *r*_*µ*_ *> r*_*s*_ (the silent phenotype is less fit), we require

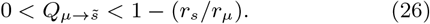

At intermediate *Q* values, the mutational clone can maintain a less-fit, yet immunogenically silent clone in place. At higher *Q* values, the model predicts a threshold consistent with Eigen’s error catastrophe [64]: if the mutator phenotype is too unstable, the *c*_*µ*_ population will be lost to its competitor.

### Stable strategies and evolutionary branching

To analyze the basic evolutionary dynamics of the trait *µ*, we link the replication function Γ(*µ*) to invasion fitness within the context of adaptive dynamics [65] as previously done in [20] to study the evolution of mutational rates. In the simplest adaptive dynamics setting, e.g. assuming frequency-independent selection in the context of the replicator equation above [59; 66], the invasion fitness of a rare mutant with trait *µ*_*m*_ in a resident population with trait *µ* can be approximated by the difference in their intrinsic replication rates: *f* (*µ*_*m*_, *µ*) *≈* Γ(*µ*_*m*_) − Γ(*µ*). Assuming mutations are rare and selection acts gradually, the direction of evolution is governed by the selection gradient, *∂f/∂µ*_*m*_ evaluated at *µ*_*m*_ = *µ*. Trait values where this gradient vanishes are candidate evolutionary singular points. At such points, directional evolution temporarily halts. Whether evolution stabilizes or diversifies around this point depends on higher-order derivatives of *f*. In the context of our work, we want to understand whether populations will halt at peaks of Γ and, more importantly, whether they will diversify at branching points produced by valleys.

To determine whether a singular point is an Evolutionarily Stable Strategy (ESS) or a branching point, we examine the second-order derivatives of *f*. A point is an ESS if small deviations by mutants reduce their invasion fitness (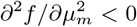 at *µ*_*m*_ = *µ*_*r*_), implying local stability against invasion. If, however, this second derivative is positive while the selection gradient vanishes, the point is a potential branching point: it is convergence stable (evolution slows down close to it) but not evolutionarily stable, allowing for divergence of nearby mutant strategies. These criteria allow us to classify the evolutionary outcomes directly from the curvature and slope of the replication function Γ(*µ*).

At *µ* = 0, going back to equations (10-11) we find that the gradient of the invasion fitness *f* is *N*_*o*_*δ*_*o*_ − *rN*_*h*_ − *N*_*a*_*δ*_*a*_, which does not vanish except from some very specific and unlikely combinations of parameter values. In the mutational tumor regime, the gradient at *µ* = 0 is positive and evolution would push tumors towards accumulating mutations and increasing *µ*. In the rest of the phase space of figure 2A, the gradient at *µ* = 0 is negative. This implies a tendency towards reducing *µ* to avoid the pressures of immune surveillance and detrimental house-keeping mutations. Because *µ* is a fraction of mutated genes, it cannot be smaller than zero by definition. Hence, *µ* = 0 in these instances is not an ESS, but rather an absorving state of the dynamics because it cannot become negative.

Instead, a branching point and an ESS can exist if there are domains in *µ* ∈ [0; 1] in which the gradient of *f* vanishes. The condition for a bimodal landscape of equations (15-16) can then be used to identify a branching point and an ESS. To establish which is which, one would compute the second derivative at this values, which can only be done numerically, and measure its sign. This can be done by visually inspecting the shape of the landscape and its first two derivatives. The first derivative can be directly inferred from Equation 11. The second derivative writes

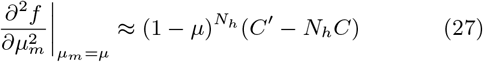

Where

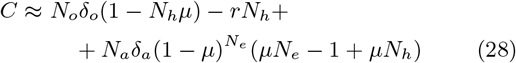

and

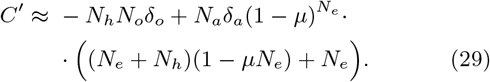

Using the results from Equations (2), (11) and (26-28), we plot in figure 5 the three functions. This allows us to easily visualize how, as expected from the simplicity of the landscape in place, the bottom valley is a branching point and the peak is an ESS of the potential evolutionary dynamics on the landscape. This brings forward the possibility that both the absorving state at *µ* = 0 and the mutational peak at *µ >* 0 are both local optima and hence cancer cells could evolve towards one or the other mutational states.

**FIG. 5.**
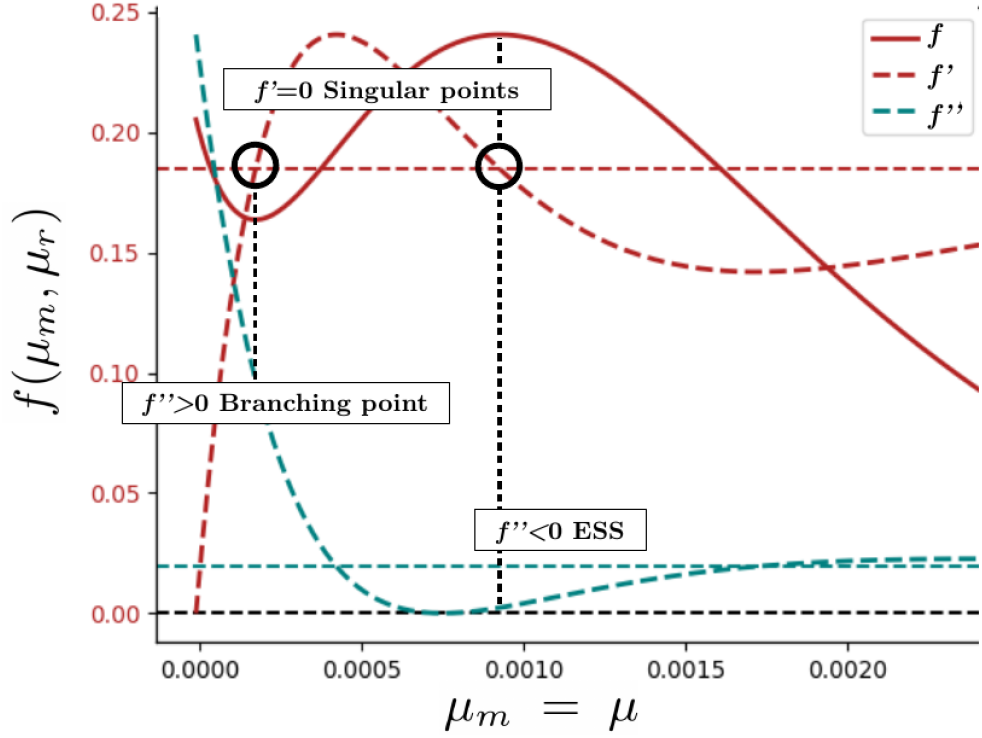
Materials and Methods : Derivatives of the evolutionary landscape *f* (*µ*_*m*_, *µ*_*r*_) *≈* Γ(*µ*_*m*_) − Γ(*µ*_*r*_). The y axis of *f* ^′^ and *f* ^′′^ are not shown as we are only interested in their sign, so that we plot their respective *y* = 0 line. This allows to observe that both points are singular points as expected where *f* ^′^ = 0, while the first is a branching point (*f* ^′′^ *>* 0) and the second is an evolutionary stable strategy (ESS, *f* ^′′^ *<* 0

### Stochastic and spatial simulations

In our simulations, we model spatial dynamics on a twodimensional lattice of fixed size (100×100), where each site is occupied by a cell of type 1 or type 2, corresponding to silent and mutational clones respectively. The aim is to explore how different spatial dynamics determine the survival time of the slow replicator silent clone inside an aggressive and mutational tumor. We explore two distinct initialization and update protocols to capture different spatial structures and dynamics. The initial spatial distribution can be random with 50% of cells of each type, or a clustering in which the slow replicator is located in an approximate disc at the center of the grid. The update scheme can be random, so that a replicating cell occupies the space of a dying cell randomly in the grid (a Moran process), or local, so that a cell can only replicate occupying the space of a nearest neighbour in the Von Neumann convention. Replication events follow a stochastic, rate-proportional (Gillespie-style) process, and simulations are run until type 1 cells go extinct or time reaches 10^7^ iterations, which explains the saturation of extinction times in figure 3C for *r*_1_ = *r*_2_. We define 30 steps between *r*_1_ = *r*_2_ = 0.2 and *r*_1_ = 0.25, and for each *r*_1_ value we perform 20 iterations and plot the mean extinction time across iterations as well as the min and max times in a shaded area.

All codes are available at:

https://github.com/GuimAguade/TMB.

